# Assessing the impact of carrier solvent and solid phase extraction blank toxicity on fish embryo testing

**DOI:** 10.64898/2025.12.01.691537

**Authors:** Jakob Pfefferle, Sarah Johann, Henner Hollert, Riccardo Massei

## Abstract

Bioassays are powerful tools to comprehensively understand the effects of complex environmental mixtures on humans and other animal species. In particular, the fish embryo test (FET) with zebrafish (*Danio rerio*) is one of the most used bioanalytical tools to test in vivo the effect of environmental extract. However, it is well known that the preparation technique of complex environmental samples for bioanalytical testing could have huge impact on the composition of exposure solutions and hence the corresponding toxicity Considering this, it is important to develop reliable environmental sample preparation procedures to obtain accurate and reproducible results in the zebrafish embryo toxicity studies. In the present study, we aimed to investigate how different experimental set-ups using a Solid Phase Extraction (SPE) extract of a complex water sample might influence the toxicity assessment using embryos of *Danio rerio*. Firstly, we assess the influence of two typical carrier solvents (methanol and dimethyl sulfoxide) for dosing environmental extracts. We further investigated the effect of increasing exposure media pH. Finally, we studied the effect of the SPE blank on lethal and sublethal effects in embryos of *Danio rerio*. Although no significant differences were observed for different solvents, we observed an adverse effect on hatching rate when testing the blank. This indicates the potential impact of background contamination in experimental setups, highlighting the need for appropriate blank correction methods. Furthermore, the differential effects observed with different pH correction methods highlight the importance of carefully selecting the appropriate pH adjustment strategy to accurately assess toxicity levels. Overall, our study enhances the knowledge base and methodology for accurate and comprehensive ecotoxicological assessments in aquatic ecosystems, aiding in the protection and conservation of these fragile environments.

## 1. Introduction

Bioanalytical tools play an important role in assessing the toxicity of environmental samples and supporting water quality monitoring (Neale et al., 2021). Together with chemical analyses, bioassays provide additional information about the combined effects of mixtures and impact of unknown chemicals present in the samples (Altenburger et al., 2019; Neale et al., 2021). The combined approach of chemical analysis and bioassays has been widely employed in characterizing various environmental samples, including matrices such as water and sediments (Hashmi et al., 2018; König et al., 2017; Massei et al., 2019; Reiter et al., 2023). *In vivo* bioassays are powerful tools to comprehensively understand the effects of complex environmental mixtures on humans and other animal species, considering essential toxicological processes such as uptake, metabolism, and excretion (Santos et al., 2018). Numerous studies have focused on utilizing the fish embryo acute toxicity (FET) test with zebrafish (*Danio rerio)* to characterize the toxicity of environmental samples. Besides the classical determination of acute lethality and morphological alterations, a wide array of toxicological endpoints providing already mechanistic understanding of underlying toxicity pathways have been established in the context of the FET (Massei et al., 2019; Perrichon et al., 2016; Rothe et al., 2021; Thellmann et al., 2015; Välitalo et al., 2017). Within those bioanalytical tools an appropriate sampling and sample preparation strategy is important to ensure meaningful toxicity assessment (Naele et al., 2021, Ribeiro et al., 2014). It is well known that the preparation technique of complex environmental samples for bioanalytical testing could have huge impact on the composition of exposure solutions and hence the corresponding toxicity (Huang et al., 2022; Kolkman et al., 2013). Solid-phase extraction (SPE) is one of the most widely used techniques to pre-enrich the sample and establish a dose-response relationship for various lethal endpoints (Neale et al., 2018; Schulze et al., 2017). Considering this, it is important to develop reliable SPE preparation procedures to obtain accurate and reproducible results in zebrafish embryo toxicity studies. Furthermore, careful selection and validation of blank toxicity is essential to ensure the reliability of environmental monitoring studies (Neale et al., 2018; Schulze et al., 2017).

In the present study, we aimed to investigate how different experimental set-ups using an SPE extract of a complex water sample might influence the toxicity assessment using embryos of *Danio rerio*. Firstly, we decided to assess the influence of different carrier solvents for dosing environmental extracts. In fact, SPE is typically performed using solvents that cannot be directly dosed in the bioassay and a changeover to compatible solvent (i.e. dimethyl sulfoxide and methanol) is necessary (Neale et al., 2021). However, different carrier solvents might modulate the solubility and bioavailability of toxic compounds, influencing their uptake and distribution within organisms. We further investigated the effect of exposure media pH on the sample toxicity since this can also strongly influence the substance uptake and toxicity (Bittner et al., 2018). In many experiments, the pH of the exposure media is not controlled through the exposure and zebrafish have a wide range of pH tolerance (Andrade et al., 2017). Finally, we studied the effect of the SPE blank on lethal and sublethal effects in embryos of *Danio rerio* and discussed potential factors linked to blank toxicity. This study provides valuable insights into the implications of sample preparation techniques on FET outcomes, contributing to improved accuracy and reliability in assessing environmental sample toxicity.

## 1. Material and methods

### 1.1 LVSPE sampling and sample preparation

An environmental sample collected from an area impacted by greenhouse activities south of Amsterdam (Netherlands) was collected in May 2018 using onsite large volume solid phase extraction (LVSPE). Shortly, 1000 L of water were extracted on site by LVSPE (Schulze et al., 2017; Välitalo et al., 2017) on a Chromabond® HRX sorbent, a hydrophobic polystyrene-divinylbenzene. The loaded HRX cartridge was freeze dried and extracted with ethyl acetate and methanol (MeOH). The extract was concentrated and recovered in 850 mL MeOH (concentration factor of 1,000). An empty cartridge was brought on field and along the sampling in order to simulate a field blank.

### 1.2 Sample preparation for the different experimental set-ups

Five aliquots of 1 mL of the extract were pipetted in 2 mL glass vials and gently evaporated under nitrogen. Two aliquots were evaporated till dryness and dissolved in MeOH and dimethyl sulfoxide (DMSO). Two further aliquots were evaporated near dryness (ca. 10 µL) and dissolved in MeOH and DMSO. All aliquots had a final concentration factor of 50,000. the fifth aliquot (completely dried) was dissolved directly in the embryo exposure medium (remineralized deionized water according to ISO 15088; 294.0 mg/L CaCl_2_ · 2H_2_O; 5.5 mg/L KCl; 123.3 mg/L, MgSO_4_ · 7H_2_O; 63.0 mg/L NaHCO_3_, pH 7-8) to a final relative enrichment factor (REF) of 100. Furthermore, one aliquot of 1 mL of the field blank was pipetted in 2 mL glass vials and gently evaporated under nitrogen. The aliquot was evaporated till dryness and dissolved in MeOH to a final concentration factor of 50,000. An overview on the different testing strategies is provided in Fig.1.

**Fig. 1.**
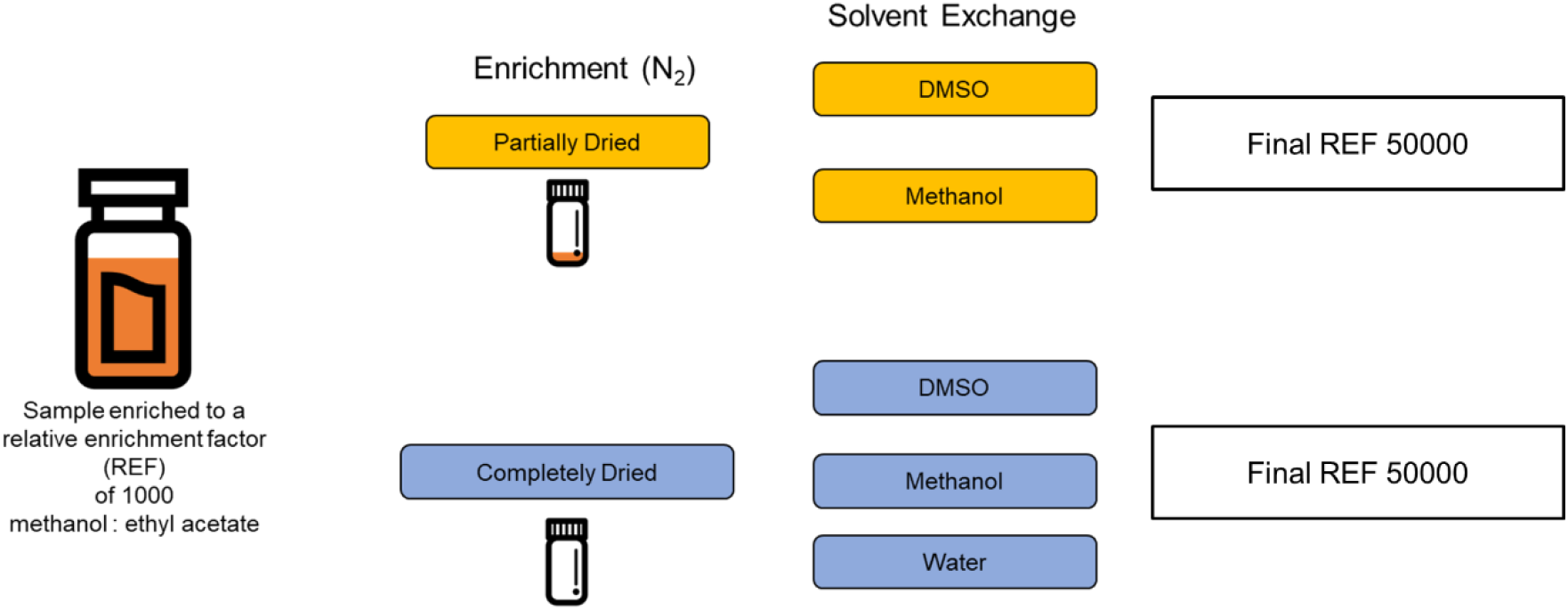
Overview of environmental sample preparation strategies after LVSPE to evaluate the impact of carrier solvent and blank toxicity for bioanalytical testing. Resulting extracts were investigated for acute embryo toxicity (FET) using zebrafish. In addition to the 5 extracts one field blank was assessed (evaporation till dryness, dissolved in MeOH, not shown). Water refers to the artificial fish embryo medium according to ISO 15088.

### 1.3 Ecotoxicological testing

#### 2.3.1 Adult fish maintenance

Adult zebrafish of the strain UFZ-OBI had been originally established from a wildtype strain purchased from a local supplier (OBI hardware store, Leipzig) and had been bred at the UFZ for more than 13 generations. Fish were kept in 14 L aquaria with 25–30 fish each with a sex distribution between female to male of 1:2. Wildtype adult zebrafish at Goethe University were kept in 167 L tanks of a recirculating system equipped with biofilter and UV-sterilization and an automatic water exchange rate of 40% per week with remineralized reverse osmosis water (remineralization with sea salt, AquaForest, Poland). For both cultures, the light–dark rhythm was 14:10 h and the water temperature was 26 ± 1 °C. Water parameters were measured frequently (pH 7–8; water hardness 2–3 mmol/L, conductivity 540–560 µS/cm, nitrate < 2.5 mg/L, nitrite < 0.025 mg/L, ammonia < 0.6 mg/L, oxygen saturation 87–91%). Within 30 min after spawning, eggs were collected using a grid covered dish and successively cleaned with aerated ISO standard dilution water (ISO-water) as specified in ISO 7346-3. Developmental stages were identified according to Kimmel et al. (1995).

#### 2.3.1 The FET with Danio rerio

The FET with zebrafish was started shortly after fertilization of the eggs (0 - 2 hours post fertilization, hpf). Fertilized eggs were separated from non-fertilized eggs after 1 h (four to eight cell stages). Non-fertilized eggs and eggs with obvious irregularities or injuries were excluded. For experiments testing the effect of solvent exchange and blank toxicity, which were performed at the UFZ Leipzig, the viable fertilized eggs were washed with ISO water and transferred to 25 mL Petri glass dishes (7 cm in diameter, VWR, Darmstadt, Germany) — glass Petri dishes were used to decrease the possibility of sorption of lipophilic substances to the test chamber. The experiments evaluating the effects of the pH switch, performed at Goethe University Frankfurt, were conducted in 96-well plates (CytoOne®, STARLAB GmbH, Hamburg, Germany). 10 viable fertilized eggs per concentration were transferred into single wells in a final exposure volume of 200 µL.

Test concentrations were calculated as relative enrichment factors (REFs) which describe how much the water sample was enriched prior to the FET and have the net units of volume extract /volume bioassay (Escher et al., 2015). All tests were performed with a final solvent concentration of 0.2 % MeOH and DMSO, respectively. For testing the effect of solvent exchange, the extract was tested in duplicates at five different REFs (10, 25, 50, 75 and 100). For testing the effect of pH switch, the extract was tested in triplicate at five different REFs (10, 25, 50, 75 and 100). Here we used buffered systems (MOPS 10 mM, HEPES 10 mM) for three different pH (6.8, 7.4, 8.2) to maintain constant pH conditions over FET duration. The pH was checked daily over the entire test period. Blank toxicity was tested at REF 20 for the LVSPE blank sample. In order to test the potential adverse effect of ammonium present in the LVSPE blank sample, which is normally higher than 8 mg/L, we further exposed zebrafish embryos to different concentration (120, 60, 30, 15, 7.5 mg/L)of ammonia (NH_3_^-^) adjusted to pH 7.4 by using either HCl (1 M)or Formic Acid (1 M). One negative (ISO-Water) and one positive control (3,4-dichloroaniline, 4 mg/L) were tested in each experiment. In total, we exposed 20 embryos with the ratio of one embryo /mL of exposure media (final volume: 10 mL).

For all experiments, exposure was started not later than 2 hpf according to OECD 236 guidelines (OECD 2013) and the tests were terminated at 96 hpf. No animal test authorization was required according to European legislation (Strähle et al. 2012). Every 24 h the embryos were checked for apical endpoints indicative of mortality (i.e. coagulation of fertilized eggs, lack of somite formation, lack of detachment of the tail-bud from the yolk sac and lack of heartbeat) and sub-lethal endpoints described in the OECD 236. Dead embryos were removed daily to avoid fungal growth. The glass Petri dishes (experiments UFZ) and well plates (experiments Goethe University) were stored in an incubator at 26±1 °C with a light-dark-cycle of 14:10 h. Experiments were counted valid if no more than 10 % of the negative control and at least 30 % of the positive control embryos showed lethal effects.

### 1.4 Data Evaluation

Percentage of mortality and (sub)lethal effects (including hatching success) were calculated for treatment and control scenarios. Afterwards, concentration-response curves were modelled with a 4-parameter non-linear regression model with variable slope. Top and Bottom were fixed at 100 and 0, respectively (Prism 9, GraphPad Software).

### 2.5 Availability of data and materials

Effect data are publicly shared on Zenodo at the addresshttps://zenodo.org/records/17777415

## 2. Results

### 2.1 Effect of carrier solvents and pH changes

The concentration-response curves obtained from our experiments showed that extracts prepared using different procedures and carrier solvents did not significantly affect the toxicity of the environmental samples (Figure 2). Extracts fully and partially re-dissolved in MeOH exhibited a mortality rate of 50 % (LC_50_) at REFs of 26 and 27, respectively. Both extracts in DMSO resulted in LC_50_ values of REF 32 (partially dried) and REF 29 (fully dried). A slight decrease in toxicity was observed for the sample re-dissolved directly in the exposure media (LC_50_: 47). Similarly, altering the pH to more acidic or alkaline levels did not appear to influence the toxicity, as the extracts exhibited comparable LC50 values after 24 (LC_50_: 23, 31 and 22 at pH 6.8, 7.4 and 8.1, respectively) and 120 (LC_50_: 20, 29 and 20 at pH 6.8, 7.4 and 8.1, respectively) hours of developmental stage (Fig. 3). Finally, sublethal effects were divergent and not consistent during the observed exposure period across different treatments.

**Fig. 2.**
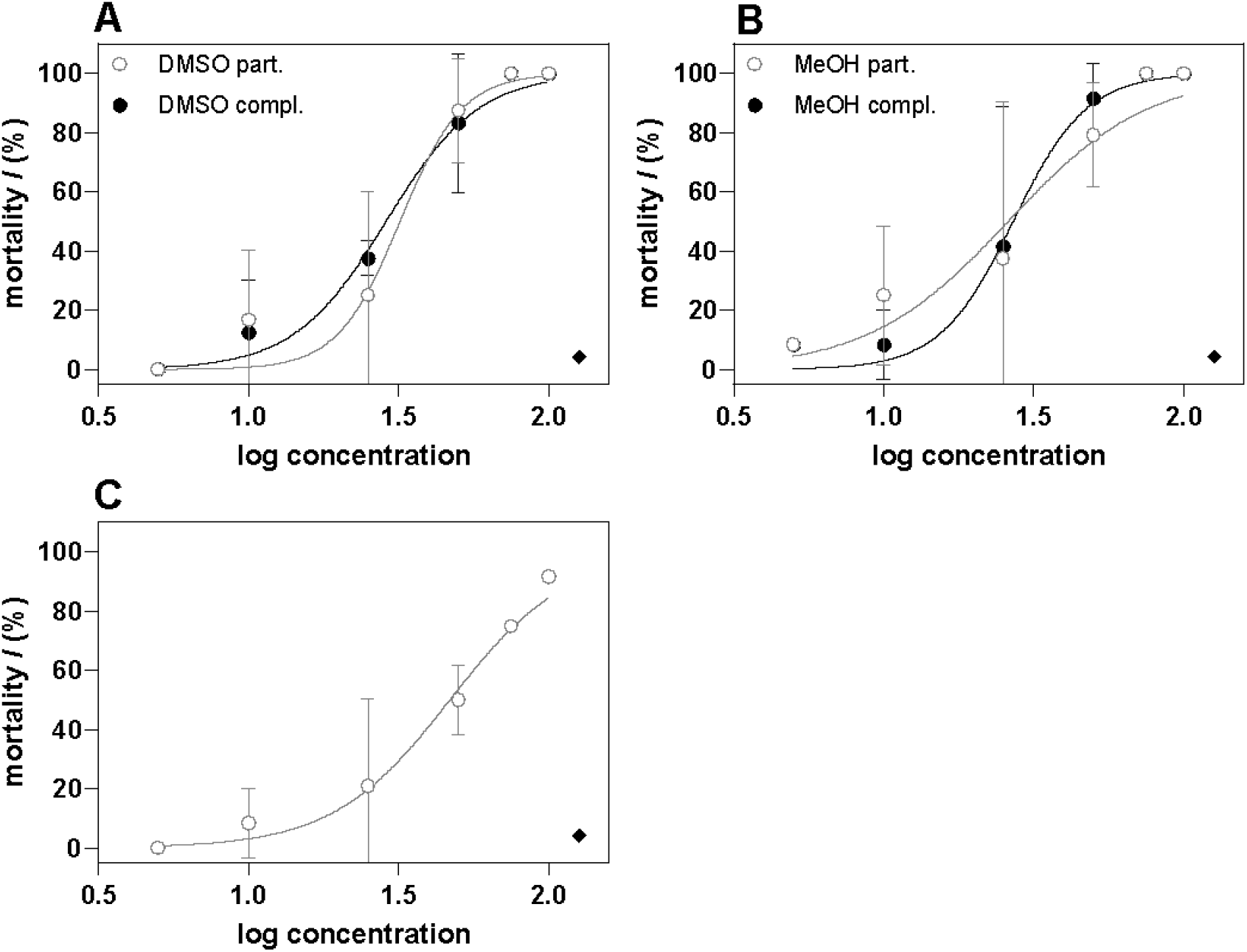
Lethal effects in zebrafish embryos (96 hpf) exposed to different sample preparation scenarios of one LVSPE extract. Extracts fully and partially dried were redissolved in DMSO (panel A), MeOH (panel B) or directly in embryo medium (water, fully dried only, panel C). Maximum solvent content 0.2 % for all treatments. n=2 presented as individual dots. Concentration-response curves were fitted using a 4-parameter non-linear regression model (top, bottom fixed at 100, 0) with variable slope using GraphPad Prism.

**Fig. 3.**
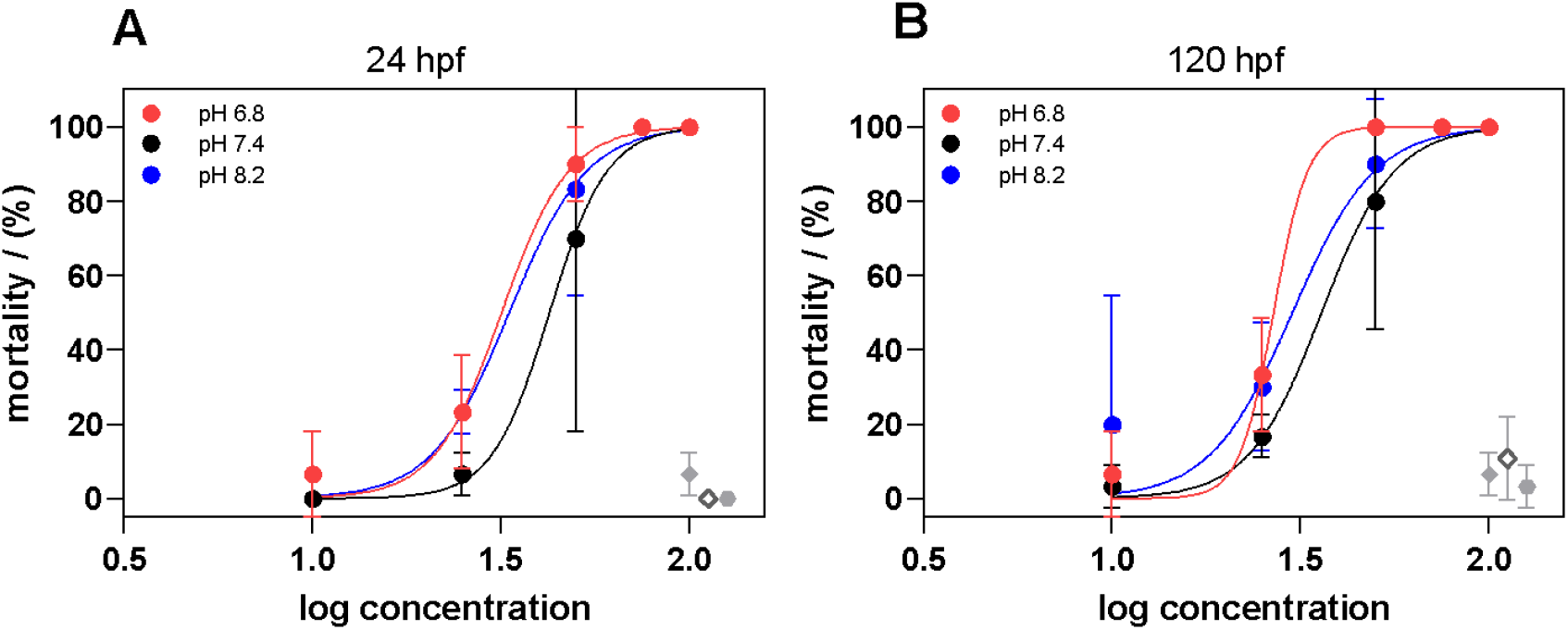
Lethal effects in zebrafish embryos (24 (panel A) and 120 (panel B) hpf) exposed to completely dried LVSPE extract redissolved in MeOH and at three different pH values Maximum solvent content was 0.2 %. n=3 presented as mean ± SD. Concentration-response curves were fitted using a 4-parameter non-linear regression model (top, bottom fixed at 100, 0) with variable slope using GraphPad Prism..

### 2.2 LVSPE blank toxicity and blank pH correction

While the LVSPE blank did not significantly affect mortality up to a REF of 20, we observed a pronounced effect on the hatching rate of the embryos (Data not shown). None of the embryos was hatched at 96 hpf. Interestingly, embryos exposed to ammonia corrected with HCl did not show any effect on mortality. However, when pH of ammonia exposure solutions was corrected with formic acid, we observed a strong increase in toxicity (LC_50_: 105 mg/L) accompanied by a decrease in the hatching rate, although the trend was not concentration-dependent (Fig. 4).

**Fig. 4.**
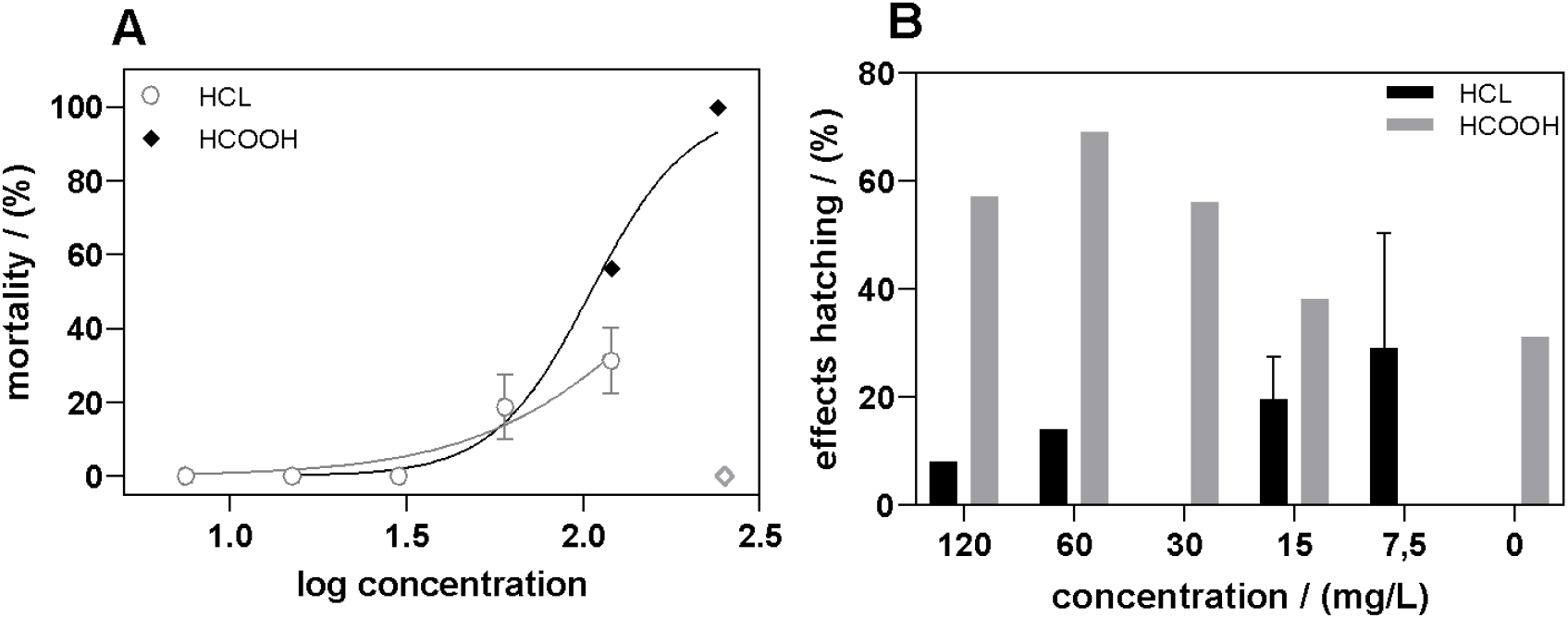
Lethal effect (A) and hatching effect (B) in zebrafish embryos (96 hpf) exposed to blank LVSPE extract (REF 20) corrected with hydrochloric acid (HCl) and formic acid (HCOOH).

## 3. Discussion

### 3.1 Carrier solvent and pH adjustment

Ensuring the use of safe and validated SPE procedures is critical for obtaining accurate and reliable bioanalytical results. Although carrier solvents and pH are well-known variables to modulate the solubility and bioavailability of toxic compounds (Bittner et al., 2018; de Koning et al., 2015; Maes et al., 2012; Spencer et al., 2018), our findings suggest that these variables might not significantly affect the overall toxicity assessment using zebrafish embryos. The concentration-response curves obtained from our experiments revealed that different preparation procedures and carrier solvents did not significantly alter the toxicity of the environmental extracts. Although there was a slight decrease in toxicity for samples re-dissolved directly in the exposure media, this difference remained within a factor of 10 and hence might be related to biological variability. However, a marginal increase in toxicity for DMSO and MeOH extracts compared to directly dissolving in medium might be related to the increased chorion permeability due to the Solvents. Kais et al. (2013) for example have shown that DMSO concentrations of ≥ 0.1 % can facilitate the uptake of chemicals in the perivitelline space. Hence, we support previous findings that independent of the selection of solvents the maximum solvent content should be as small as possible for a realistic hazards assessment. Regarding the choice of solvent, the results demonstrate no significant differences between DMSO and MeOH. DMSO is the most commonly used solvent in bioassays, however it is known to interfere during mass spectrometric analysis (Kolkmann et al., 2013) and alters, e.g. chorion permeability. In contrast, MeOH is a suitable solvent also for the application in chemical analysis. MeOH can also be used as a solvent in bioassays, indicating its suitability for a combined approach of bioassays and chemical analysis. However, it has to be considered that MeOH is a volatile solvent meaning that the vial weight of extracts has to be continuously monitored and adjusted to keep the correct concentration/enrichment factor. DMSO is still being discussed as the solvent of choice especially for condensed matrices such as sediments (Naele et al., 2021). In contrast, MeOH is a suitable solvent also for the application in chemical analysis.

Additionally, altering the pH levels to more acidic or alkaline conditions did not appear to have a substantial influence on the toxicity, as evidenced by comparable LC_50_ values observed at 24 and 96 hours of exposure..

### 4.2 LVSPE blank toxicity and the impact of acid selection for pH adjustment

The elution process of the LVSPE cartridges is typically divided in four different steps, where the last two include the pH adjustment with MeOH/NH3 and MeOH/HCOOH. For every elution passage, a blank control is included to account for potential errors or contaminations during the process. In the past, these blanks often showed adverse effects in biotests. We hypothesized that this might be due to toxicity caused by ammonia or its reaction with acids utilized for pH adjustment. To study this impact, we exposed fish embryos to a LVSPE blank sample and to a dilution series of ammonia (NH_3_) adjusted to neutral pH (7.4) with (a) hydrochloric acid (HCl) or (b) formic acid (HCOOH). Our findings reveal that while the LVSPE blank did not significantly affect mortality, it fully inhibited hatching. Studying the potential impact of NH_3_/HCOOH/HCl on this hatching delay, we observed that HCl did not exhibit any significant effect while formic acid increased the toxicity including a decrease in the hatching rate. The difference might likely be explained by the chemical interaction of NH3 with either HCl or HCOOH. In an aqueous solution, the equilibrium of NH_3_/NH_4_ ^+^ is shifted towards one of both constituents depending on the pH, temperature and the acid used (Fu and Tian, 2013). Ammonium (NH_4_ ^+^) is not toxic to fish, while ammonia is known to cause strong adverse lethal and sublethal effects in aquatic organisms and fish in particular ((Kır et al., 2019; Luo et al., 2016; Randall and Tsui, 2002; Wang et al., 2017). For zebrafish a LC_50_ of 2.07 mg NH_3_/L was reported (Mariz Jr. et al., 2023)

At the tested pH of 7.4 and 26°C, an approximate calculated 1.5 % of the total ammonia is un-ionized NH_3_ based on calculations of Emerson et al. (1975), which indicated an overall low toxicity due to the shift towards the non-toxic NH_4_ ^+^. Importantly, HCl is a strong acid (pKa -7, Cotton andWilkinson 1974) while HCOOH is a rather weak acid (pKa 3.77, Brown et al., 1955). This indicates that under the established conditions HCl is fully dissociated into H+ and Cl-, both non-toxic ions for fish embryos, whereas for HCOOH the acid itself as well as ammonium formate (HCOONH_4_) will be present due to limited dissociation and reaction with NH3. HCOONH4 has been reported to cause lethal effects in fish in the mg/L range (e.g. 175 mg/L: Dowden and Bennett, 1965). Another aspect that could impact the overall higher toxicity within the NH3/HCOOH experiment is that HCOOH can be metabolized to formaldehyde (CH_2_O).

## 5. Conclusion

Though no significant differences were observed for different solvents and pH values, researchers must carefully consider the choice and concentration of carrier solvents when conducting experiments with zebrafish embryos to minimize potential confounding effects and ensure the accuracy and reliability of their findings. These results emphasize the significance of considering LVSPE blanks and pH correction in ecotoxicological studies using zebrafish embryos as model organisms. The adverse effect on the hatching rate indicates the potential impact of background contamination in experimental setups, highlighting the need for appropriate blank correction methods. Furthermore, the differential effects observed with different pH correction methods highlight the importance of carefully selecting the appropriate pH adjustment strategy to accurately assess toxicity levels. Based on the present findings we recommend avoiding NH3 and HCOOH within elution steps of LVSPE.

In conclusion, our results contribute to the understanding of the reliability and robustness of using zebrafish embryos for toxicity evaluations of environmental samples. It is important to consider these factors when designing and conducting ecotoxicological studies, as they provide valuable insights into the potential risks associated with chemical exposures. Future research should focus on further investigating the mechanisms underlying the observed toxicity patterns and exploring additional endpoints to assess sublethal effects. Overall, our study enhances the knowledge base and methodology for accurate and comprehensive ecotoxicological assessments in aquatic ecosystems, aiding in the protection and conservation of these fragile environments.

## 6. Acknowledgement and fundings

The authors would like to acknowledge the contribution of the NORMAN for funding (https://www.norman-network.net/) and in particular the participants in the WG-2 Bioassay

## 7. Declaration on LLM usage for grammar correction

In the present manuscript, LLM (ChatGPT 4.0) was used for grammar revision, and this usage is disclosed in accordance with institutional guidelines.

## References

Altenburger, R., Brack, W., Burgess, R.M., Busch, W., Escher, B.I., Focks, A., Mark Hewitt, L., Jacobsen, B.N., de Alda, M.L., Ait-Aissa, S., 2019. Future water quality monitoring: improving the balance between exposure and toxicity assessments of real-world pollutant mixtures. Environmental Sciences Europe 31, 1–17.

Andrade, T.S., Henriques, J.F., Almeida, A.R., Soares, A.M.V.M., Scholz, S., Domingues, I., 2017. Zebrafish embryo tolerance to environmental stress factors—Concentration–dose response analysis of oxygen limitation, pH, and UV-light irradiation. Environmental Toxicology and Chemistry 36, 682–690. 10.1002/etc.3579.

Bittner, L., Teixido, E., Seiwert, B., Escher, B.I., Klüver, N., 2018. Influence of pH on the uptake and toxicity of β-blockers in embryos of zebrafish, Danio rerio. Aquatic Toxicology 201, 129–137.

Brown, H.C., McDaniel, D.H., Häfliger, O., 1955. Determination of Organic Structures by Physical Methods - Dissociation constants. Academic Press.

de Koning, C., Beekhuijzen, M., Tobor-Kapłon, M., de Vries-Buitenweg, S., Schoutsen, D., Leeijen, N., van de Waart, B., Emmen, H., 2015. Visualizing compound distribution during zebrafish embryo development: the effects of lipophilicity and DMSO. Birth Defects Research Part B: Developmental and Reproductive Toxicology 104, 253–272.

Dowden, B.F., Bennett, H.J., 1965. Toxicity of Selected Chemicals to Certain Animals. Journal (Water Pollution Control Federation) 37, 1308–1316.

Emerson, K., Russo, R.C., Lund, R.E., Thurston, R.V., 1975. Aqueous Ammonia Equilibrium Calculations: Effect of pH and Temperature. J. Fish. Res. Bd. Can. 32, 2379–2383. 10.1139/f75-274.

Fu, C.-F., Tian, S.X., 2013. Thermodynamics of Ammonia and Ammonium Ion at the Aqueous Solution–Air Interfaces. J. Phys. Chem. C 117, 13011–13020. 10.1021/jp312110w.

Hashmi, M.A.K., Escher, B.I., Krauss, M., Teodorovic, I., Brack, W., 2018. Effect-directed analysis (EDA) of Danube River water sample receiving untreated municipal wastewater from Novi Sad, Serbia. Science of the total environment 624, 1072–1081.

Huang, S., Fan, M., Wawryk, N., Qiu, J., Yang, X., Zhu, F., Ouyang, G., Li, X.-F., 2022. Recent advances in sampling and sample preparation for effect-directed environmental analysis. TrAC Trends in Analytical Chemistry 154, 116654. 10.1016/j.trac.2022.116654.

Kais, B., Schneider, K.E., Keiter, S., Henn, K., Ackermann, C., Braunbeck, T., 2013. DMSO modifies the permeability of the zebrafish (Danio rerio) chorion-implications for the fish embryo test (FET). Aquatic toxicology 140, 229–238.

Kimmel, C.B., Ballard, W.W., Kimmel, S.R., Ullmann, B., Schilling, T.F., 1995. Stages of embryonic development of the zebrafish. Developmental dynamics 203, 253–310.

Kr, M., Sunar, M.C., Gök, M.G., 2019. Acute ammonia toxicity and the interactive effects of ammonia and salinity on the standard metabolism of European sea bass (Dicentrarchus labrax). Aquaculture 511, 734273. 10.1016/j.aquaculture.2019.734273.

Kolkman, A., Schriks, M., Brand, W., Bäuerlein, P.S., van der Kooi, M.M.E., van Doorn, R.H., Emke, E., Reus, A.A., van der Linden, S.C., de Voogt, P., Heringa, M.B., 2013. Sample preparation for combined chemical analysis and in vitro bioassay application in water quality assessment. Environmental Toxicology and Pharmacology 36, 1291–1303. 10.1016/j.etap.2013.10.009

König, M., Escher, B.I., Neale, P.A., Krauss, M., Hilscherová, K., Novák, J., Teodorović, I., Schulze, T., Seidensticker, S., Hashmi, M.A.K., 2017. Impact of untreated wastewater on a major European river evaluated with a combination of in vitro bioassays and chemical analysis. Environmental Pollution 220, 1220–1230.

Luo, S., Wu, B., Xiong, X., Wang, J., 2016. Short-term toxicity of ammonia, nitrite, and nitrate to early life stages of the rare minnow (Gobiocypris rarus). Environmental Toxicology and Chemistry 35, 1422–1427. 10.1002/etc.3283.

Maes, J., Verlooy, L., Buenafe, O.E., De Witte, P.A., Esguerra, C.V., Crawford, A.D., 2012. Evaluation of 14 organic solvents and carriers for screening applications in zebrafish embryos and larvae.

Mariz Jr., C.F., Melo Alves, M.K. de, Santos, S.M.V. dos, Alves, R.N., Carvalho, P.S.M., 2023. Lethal and Sublethal Toxicity of Un-Ionized Ammonia to Early-Life Stages of Danio rerio. Zebrafish 20, 67–76. 10.1089/zeb.2022.0064

Massei, R., Hollert, H., Krauss, M., Von Tümpling, W., Weidauer, C., Haglund, P., Küster, E., Gallampois, C., Tysklind, M., Brack, W., 2019. Toxicity and neurotoxicity profiling of contaminated sediments from Gulf of Bothnia (Sweden): a multi-endpoint assay with Zebrafish embryos. Environmental Sciences Europe 31, 1–12.

Neale, P., Leusch, F., Escher, B., 2021. Bioanalytical tools in water quality assessment. IWA publishing.

Neale, P.A., Brack, W., Aït-Aïssa, S., Busch, W., Hollender, J., Krauss, M., Maillot-Maréchal, E., Munz, N.A., Schlichting, R., Schulze, T., 2018. Solid-phase extraction as sample preparation of water samples for cell-based and other in vitro bioassays. Environmental Science: Processes & Impacts 20, 493–504.

Perrichon, P., Le Menach, K., Akcha, F., Cachot, J., Budzinski, H., Bustamante, P., 2016. Toxicity assessment of water-accommodated fractions from two different oils using a zebrafish (Danio rerio) embryo-larval bioassay with a multilevel approach. Science of the Total Environment 568, 952–966.

Randall, D.J., Tsui, T.K.N., 2002. Ammonia toxicity in fish. Marine Pollution Bulletin 45, 17–23. 10.1016/S0025-326X(02)00227-8.

Reiter, E.B., Escher, B.I., Rojo-Nieto, E., Nolte, H., Siebert, U., Jahnke, A., 2023. Characterizing the marine mammal exposome by iceberg modeling, linking chemical analysis and in vitro bioassays. Environmental Science: Processes & Impacts.

Ribeiro, C., Ribeiro, A.R., Maia, A.S., Gonçalves, V.M.F., Tiritan, M.E., 2014. New Trends in Sample Preparation Techniques for Environmental Analysis. Critical Reviews in Analytical Chemistry 44, 142–185. 10.1080/10408347.2013.833850.

Rothe, L.E., Botha, T.L., Feld, C.K., Weyand, M., Zimmermann, S., Smit, N.J., Wepener, V., Sures, B., 2021. Effects of conventionally-treated and ozonated wastewater on mortality, physiology, body length, and behavior of embryonic and larval zebrafish (Danio rerio). Environmental Pollution 286, 117241.

Santos, D., Vieira, R., Luzio, A., Félix, L., 2018. Zebrafish early life stages for toxicological screening: insights from molecular and biochemical markers, Advances in molecular toxicology. Elsevier, pp. 151–179.

Schulze, T., Ahel, M., Ahlheim, J., Aït-Aïssa, S., Brion, F., Di Paolo, C., Froment, J., Hidasi, A.O., Hollender, J., Hollert, H., 2017. Assessment of a novel device for onsite integrative large-volume solid phase extraction of water samples to enable a comprehensive chemical and effect-based analysis. Science of the Total Environment 581, 350–358.

Spencer, H., Wahome, J., Haasch, M., 2018. Toxicity evaluation of acrylamide on the early life stages of the zebrafish embryos (Danio rerio). Journal of Environmental Protection 9, 1082.

Thellmann, P., Köhler, H.-R., Rößler, A., Scheurer, M., Schwarz, S., Vogel, H.-J., Triebskorn, R., 2015. Fish embryo tests with Danio rerio as a tool to evaluate surface water and sediment quality in rivers influenced by wastewater treatment plants using different treatment technologies. Environmental Science and Pollution Research 22, 16405–16416.

Välitalo, P., Massei, R., Heiskanen, I., Behnisch, P., Brack, W., Tindall, A.J., Du Pasquier, D., Küster, E., Mikola, A., Schulze, T., 2017. Effect-based assessment of toxicity removal during wastewater treatment. Water research 126, 153–163.

Wang, H.-J., Xiao, X.-C., Wang, H.-Z., Li, Y., Yu, Q., Liang, X.-M., Feng, W.-S., Shao, J.-C., Rybicki, M., Jungmann, D., Jeppesen, E., 2017. Effects of high ammonia concentrations on three cyprinid fish: Acute and whole-ecosystem chronic tests. Science of The Total Environment 598, 900–909. 10.1016/j.scitotenv.2017.04.070.

Wilkinson, G., Cotton, A., 1974. Anorganische Chemie, 3. Auflage. ed.

